# Long-range sensitization in the vertebrate retina and human perception

**DOI:** 10.1101/492033

**Authors:** Benjamin N. Naecker, Stephen A. Baccus

## Abstract

Sensory stimuli have extensive correlations, which should be exploited by an efficient detector of specific sensory features according to theoretical principles. Here we show that visual stimulation far outside the classical receptive field center causes ganglion cells to subsequently increase their sensitivity to local stimuli in a process we term long-range sensitization. This increased sensitivity improves detection and discrimination of weak stimuli and exhibits pattern specificity, increasing sensitivity more when peripheral and central stimuli share spatial frequency statistics. This process requires input from wide-field, nonlinear inhibitory amacrine cells, supporting a simple circuit model that reproduces sensitization. We further show that long-range sensitization parallels a novel perceptual effect in humans, in which surround stimulation subsequently improves discrimination of a small central stimulus. We conclude that the retina uses long-range statistics of the visual world to better encode local visual features, and that such improved encoding may play a role in perception.

---

The natural visual world is highly structured. Nearby regions in natural scenes tend to have similar structure^1^, and most objects move slowly or not at all^2^. These properties generate a visual world in which information about a single location at a single point in time provides strong statistical expectations about the features of nearby visual input.

Theories of efficient coding have proposed that an important function of the retinal circuit is to remove redundancy created by such correlations^3–6^. However, a large and growing body of recent work suggests that the ganglion cells, the output of the retina, act as a set of detection channels, each tuned to specific and ethologically-relevant visual features ^7–10^.

In this framework, classical theories of signal detection^11^ suggest that ganglion cells should use, rather than remove, the statistical regularities of the natural world to better detect the feature to which they are tuned. The presence of a visual feature in one location of the visual world provides evidence for its existence elsewhere, which a detector striving for optimality ought to exploit.

Previous work has shown that a subset of ganglion cells do indeed use statistics of their inputs to maintain a statistical estimate of the likely inputs over a local spatial region (<200μ*m*) for a short time ^12,13^ called local sensitization. Yet the statistical structure of natural visual inputs contains correlations over far larger spatial scales. Furthermore, the proposed mechanism for sensitization relies on adaptation in parallel excitatory and inhibitory pathways, a circuit motif which is likely to be found throughout the retina.

Here we report that visual stimulation in surrounding regions far outside the receptive field center causes a persistent increase in sensitivity to weak central stimuli. This increase in local sensitivity is larger when the local and distal inputs share visual features than when they do not, a simple form of pattern-specific statistical feature detection.

To explore the cellular and circuit mechanisms underlying long-range sensitization, we pharmacologically manipulated retinal circuitry, showing that both wide-field nonlinear and narrow-field linear inhibition are required for long-range sensitization. We propose a simple circuit model of long-range sensitization that uses well-understood retinal components as its building blocks.

To demonstrate that this computation is relevant to perception, we further show that long-range sensitization parallels a novel perceptual effect in humans, in which strong stimulation of a surrounding region improves the detection of and sensitivity to a small central stimulus. This establishes that the computation of long-range sensitization is important for the detection and processing of ethologically-relevant visual features, and that the hypothesized neural circuit provides a simple motif which may be used to implement statistical inference about a wide array of features in the visual world.

## Results

### Surround stimulation modulates ganglion cell responses

To investigate the influence of peripheral stimuli on local feature sensitivity, we presented a stimulus in which the peripheral regions of the retina were strongly stimulated, after which local ganglion cell sensitivity to could be assessed. Spatially, the stimulus consisted of a central region (500 μm), which covered the receptive field centers of ganglion cells, and a much larger surrounding region (~6 mm), which covered the entire retina. Both regions contained alternating dark and light bars in a grating pattern, with a spatial period of 100 μm (Fig. 1A). We recorded the electrical activity of ganglion cells responding to this stimulus with a multielectrode array (Fig. 1A, B). Each spatial region reversed its contrast in a square wave pattern, at a temporal period of 400ms (Fig. 1C and D).

To test the effects of strong stimulation, each trial began with an adapting period (0 to 5.2s), during which the surround contrast was high and the center contrast low (100% and 3-15% Michelson contrast) (Fig. 1A). After the adapting period, the contrast of the surround region was reduced to match that of the center (Fig. 1B, D). This probe period lasted the remaining 10s of each trial.

To characterize these effects of strong surround stimulation, we investigated ganglion cell responses in the initial portion of the probe period (*L*_*early*_, 0.5 to 2.5s after contrast switch), and during the final segment of the trial (*L*_*late*_, final 3s of trial), which provides a near steady-state condition against which sensitivity during *L*_*early*_ may be compared. After the switch from adapting to probe period, many cells showed increases in firing rate relative to the baseline steady-state level (Fig. 1E). This increased responsiveness during *L*_*early*_ decayed over the following several seconds for most cells, such that overall responsiveness was much lower during *L*_*late*_. To quantify whether these changes in firing rate reflect changes in the sensitivity of each cell, we estimated the contrast response function of each cell as the mean firing rate at each contrast separately for the time intervals *L*_*early*_ or *L*_*late*_ (Fig. 1F). We observed a large increase in the sensitivity to contrast in *L*_*early*_ as compared to *L*_*late*_. The change in slope during *L*_*early*_ compared to *L*_*late*_ was 4.57 ± 1.04 (mean ± s.e.m., p=5.85^−5^, n = 50). Therefore, strong stimulation in the surround region, far outside the receptive field center, subsequently increases the local sensitivity of ganglion cells.

**Figure 1.**
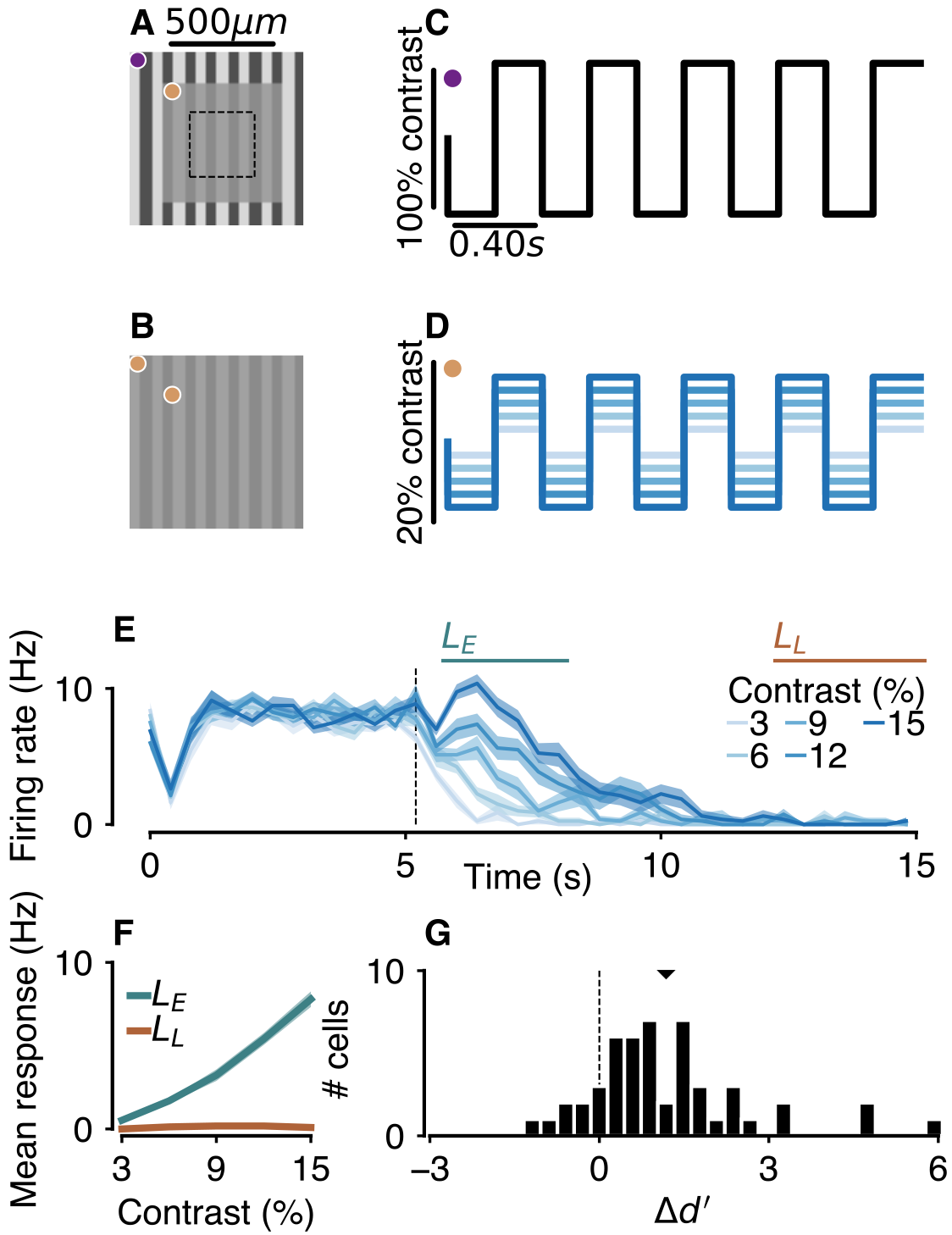
Surround stimulation increases local stimulus discriminability. Spatial (A and B) and temporal (C and D) parameters of the stimulus to probe sensitization. (A) The stimulus consisted of a central region (500μm), centered over the multielectrode array (dashed black lines), and a much larger surround region (~6mm), both containing square-wave gratings with a 100μm spatial period. The gratings in both regions reversed contrast with a temporal period of 0.40s (C and D). Each trial began with an adapter (5.2s), in which the contrast of the surround grating was 100%, while the center grating had one of several low contrasts (3-20%). During the remaining 10s of each trial, the probe, the surround grating reduced contrast to match that of the center. Purple and yellow circles in C and D indicate the temporal profile of the region marked with the corresponding color in A and B. Many trials of a single contrast were repeated, and interleaved in blocks, with a total of 20 trials of each contrast. (E) Peristimulus time histogram (PSTH) for one example cell, showing the firing rate as a function of time in each trial. Lines indicate average across 20 trials of each center contrast, and shaded areas indicate mean ±1 s.e.m. Black vertical line indicates the switch from adapter to probe periods. Blue and orange lines indicate periods of analysis, *L*_*early*_ = 5.7-8.2s and *L*_*late*_ = 12.215.2s. (F) Average firing rate of cell in (E) as a function of center contrast. Lines show average across 20 trials, and shaded areas indicate mean ±1 s.e.m. (G) Change in discriminability (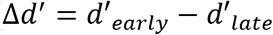 see Eq. 1) over 50 recorded RGCs, computed from response curves for each cell as in (F). Black dotted line is at zero, which indicates no change in discriminability between *L*_*early*_ and *L*_*late*_. Black arrow indicates population mean, 1.18 (s.e.m. = 0.20), significantly above 0 (two-sided *t*-test, p = 3.075^−7^).

### Surround stimulation improves local stimulus discriminability

The observed increases in firing as a function of contrast may allow similar stimuli to be more easily discriminated. However, if the response function increased so much as to saturate, then similar stimuli would be less discriminable. Additionally, increases in response variability may adversely impact discriminability, independent of changes in the mean response. A measure which takes both the mean response to similar stimuli as well as the response variability is the discriminability, *d*′, defined here as:

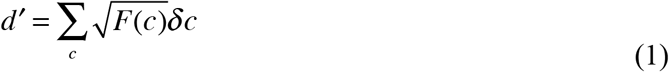

where *δc* is the contrast step between adjacent contrasts, and *F*(*c*) is the Fisher information:

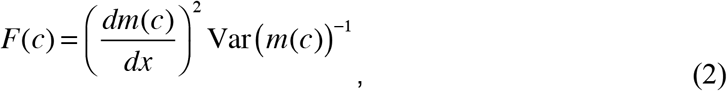

where Var(*m*(*c*)) is the variance of the contrast response function computed across trials. The Fisher information quantifies the separation (local slope) between responses generated by stimuli of similar contrast, normalized by the variability of those responses, and sets an upper bound on the accuracy of any unbiased decoding scheme^14^.

The effect of strong surround stimulation on *d*′ can be quantified by the change in discriminability Δ*d*′ between the relevant time periods, *L*_*early*_ and *L*_*late*_, which is the difference in discriminability immediately after the contrast switch compared to the near steady-state condition. Fig. 1G shows the distribution of Δ*d*′ over 50 recorded ganglion cells. The average change in discriminability over all cells was significantly shifted above zero (mean ± 1 s.e.m., 1.18 ± 0.20, two-sided *t*-test, p=3.08^−7^), indicating higher discriminability during *L*_*early*_ than *L*_*late*_. Thus, strong surround stimulation subsequently increases the sensitivity of ganglion cells.

### Long-range sensitization is specific to spatial patterns

The preceding results show that strong stimulation in regions peripheral to a ganglion cell can drive changes in the cell’s responses to central stimuli, and that these responses support better discrimination of similar stimuli. However, ganglion cells acting as feature detectors should use not only the contrast information of the surrounding visual input, but also the precise statistics of that input to better detect local features. Because textures tend to be similar across space, changes in ganglion cell sensitivity might show some form of pattern-specificity, such that the statistics of the surround enable better discrimination of local stimuli with similar statistics.

To test the pattern-specificity of long-range sensitization, we modified the stimulus in Fig. 1 to include stimuli in which the spatial correlations of the center and surround vary. In this stimulus, shown in Fig. 2A-D, both the center and surround contained either the same spatial grating pattern described above, or spatially uniform illumination, yielding four combinations. The temporal pattern of the stimulus was identical to that shown in Fig. 1C and D.

**Figure 2.**
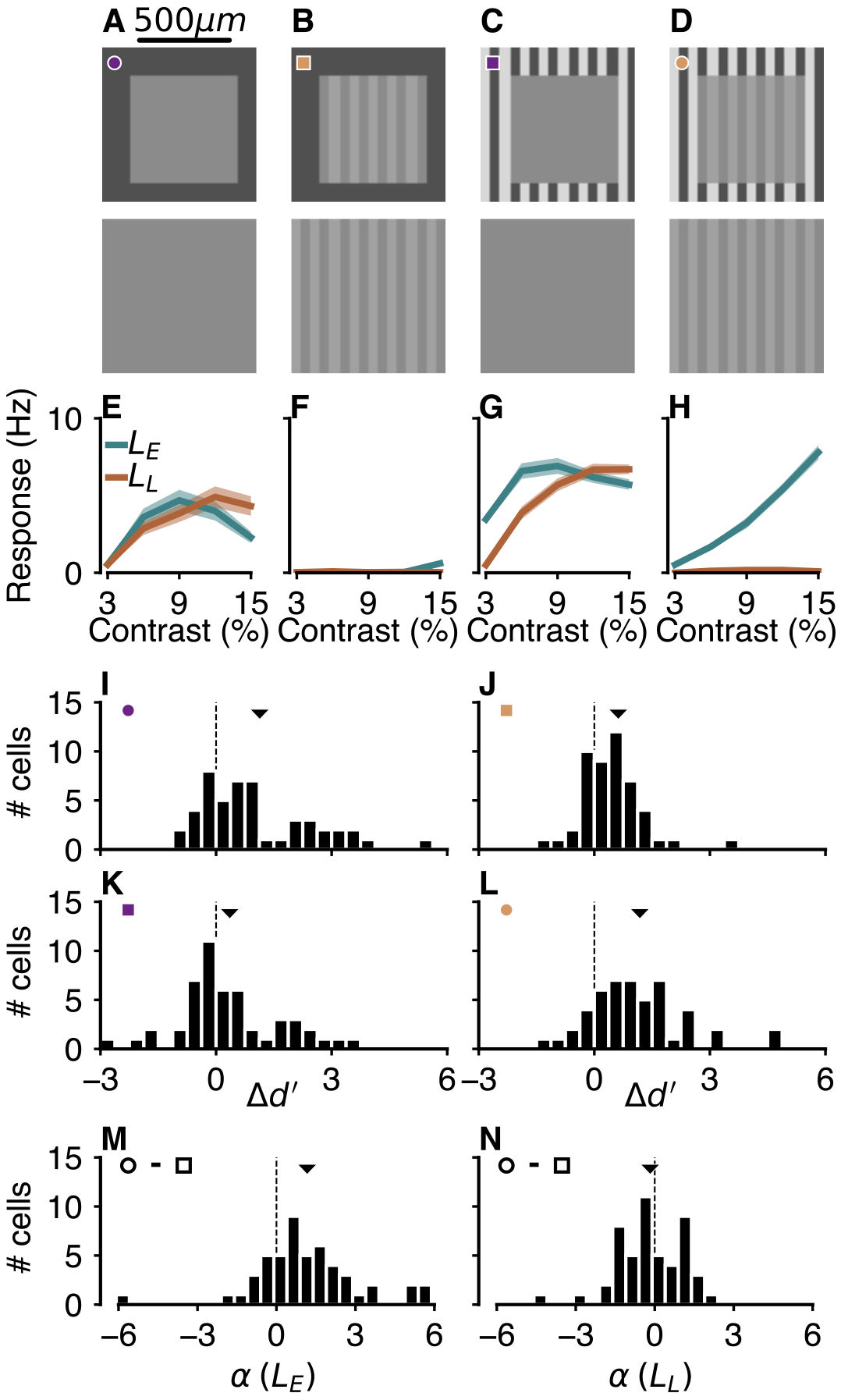
Long-range sensitization is specific to spatial pattern. (A-D) Spatial arrangement of the stimulus during the adapter (top) and probe (bottom) periods. The surround and center stimuli contained either uniform illumination or alternating grating patterns. The size of the center and surround regions, the trial durations, and the manner in which the stimuli flickered over time were all identical to those in Figure 1. (E-H) Response curves during *L*_*early*_ (green) and *L*_*late*_ time periods showing mean firing rate as a function of contrast, for one example cell, in each stimulus condition in (A-D), respectively. Shaded areas show ± 1 s.e.m. (I-L) The distribution of Δ*d*′ between *L*_*early*_ and *L*_*late*_ (see Eq. 1) over all recorded cells, with the mean of each distribution indicated by the above arrow. The stimulus condition from which the distributions were derived are indicated by the colored symbols in the top left, which match the symbols in (A-D). The mean in each panel is indicated by the black arrow. Mean ± 1 s.e.m. and p-values for two-sided *t*-tests comparing the mean to zero (indicating no change in *d*′) are: I: (1.13 ± 0.22, 7.27^−6^); J: (0.62 ± 0.19, 0.0023); K: (0.35 ± 0.18, 0.060); L: (1.18 ± 0.200, 3.075^−7^). (M and N) Pattern-specificity indices *α* during *L*_*early*_ (M) and *L*_*late*_ (N). *α* compares *d*′ during stimuli in which center and surround patterns were the same (conditions marked with circles) with those in which the patterns were different (conditions marked with squares). Values greater than zero indicate preference for same over different patterns, while values less than zero indicate preference for different over same patterns. Mean ± 1 s.e.m. and p-values for two-sided *t*-tests comparing the means to zero (indicating no preference for pattern) are: M: (1.166 ± 0.281, 0.00013); N: (−0.17 ± 0.18, 0.342).

Fig. 2A-D shows the adapter (top) and probe stimulus (bottom) for each of the four stimulus conditions. In addition to a reduction in contrast, in the probe period the surround region also changed its pattern to match that of the center (bottom row). This was done to avoid measuring direct sensitivity to the adapting pattern during the probe period. Fig. 2E-H shows the contrast response functions of an example cell in each stimulus condition. As above, we estimated the discriminability from the response curves as *d*′ (see Eq. 1) and compared this quantity in *L*_*early*_ to that in *L*_*late*_ (Fig. 2I-L). In each condition, the distribution is significantly shifted above zero, indicating that discriminability was higher during *L*_*early*_ than *L*_*late*_. (See figure legend for summary statistics.) Strong stimulation in the surround thus increases local stimulus discriminability in all cases, regardless of the correlation of the surround and center regions.

However, improvements in discriminability were larger when the surround and center correlations are the same (Fig. 2I, L), compared to when they are different (Fig. 2J, K). To measure this pattern-specificity, we computed the degree to which changes in stimulus discriminability favored trials in which the center and surround contained the same spatial structure, whether correlated or anti-correlated as:

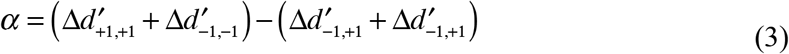

A cell with similar discriminability changes on all trials, regardless of the relationship between the surround and center stimuli, will have *α* ≈ 0. Values less than zero indicate higher discriminability on trials in which the stimuli in surround and center contained different spatial patterns, and values greater than zero indicate higher discriminability on trials in which the probe and adapter stimuli shared the same type of spatial correlation.

On average, indices of pattern specificity during *L*_*early*_ were significantly greater than zero (mean ± 1 s.e.m., 1.166 ± 0.281, *p*=1.31^−4^), whereas those during *L*_*late*_ were not (−0.17 ± 0.18 *p*=0.34). This suggests that ganglion cells use not only contrast but the higher-order correlation structure of the global visual scene to better detect stimuli in their local receptive field.

### Sensitivity changes in both the surround and center

Because the probe stimulus of Fig. 2 covered both the central and peripheral region, it cannot be determined whether the peripheral stimulus changes sensitivity to the central stimulus or the peripheral stimulus. To distinguish these possibilities, we designed a stimulus in which the center and surround regions were independently probed, shown in Fig. 3. To independently probe the surround and center region, the stimulus was segregated into three conditions (Fig. 2A). In the “Full-field” condition (top row), the stimulus was present in both the surround and center (identical to previous stimuli). The middle row shows the “Surround-only” condition, in which the surround region contained the same stimulus, but the center region was presented with uniform grey illumination throughout the entire trial duration. In the “Center-only” condition, the surround and center components were both visible during the adapter period (left), but only the center component was visible during the probe (right). Changes in discriminability in this case are due solely to modulations of excitation in the center region. We found that discriminability changed in all three conditions, indicating that sensitivity in both the central and peripheral regions was increased by peripheral stimulation (Fig. 3B, mean ± 1 s.e.m, Full-field: 1.03 ± 0.17, p=1.08^−7^; Surround-only: 0.58 ± 0.17, p=0.0014; Center-only: 0.68 ± 0.28, p=0.016).

### Influence of narrow and wide-field inhibition on sensitization

**Figure 3.**
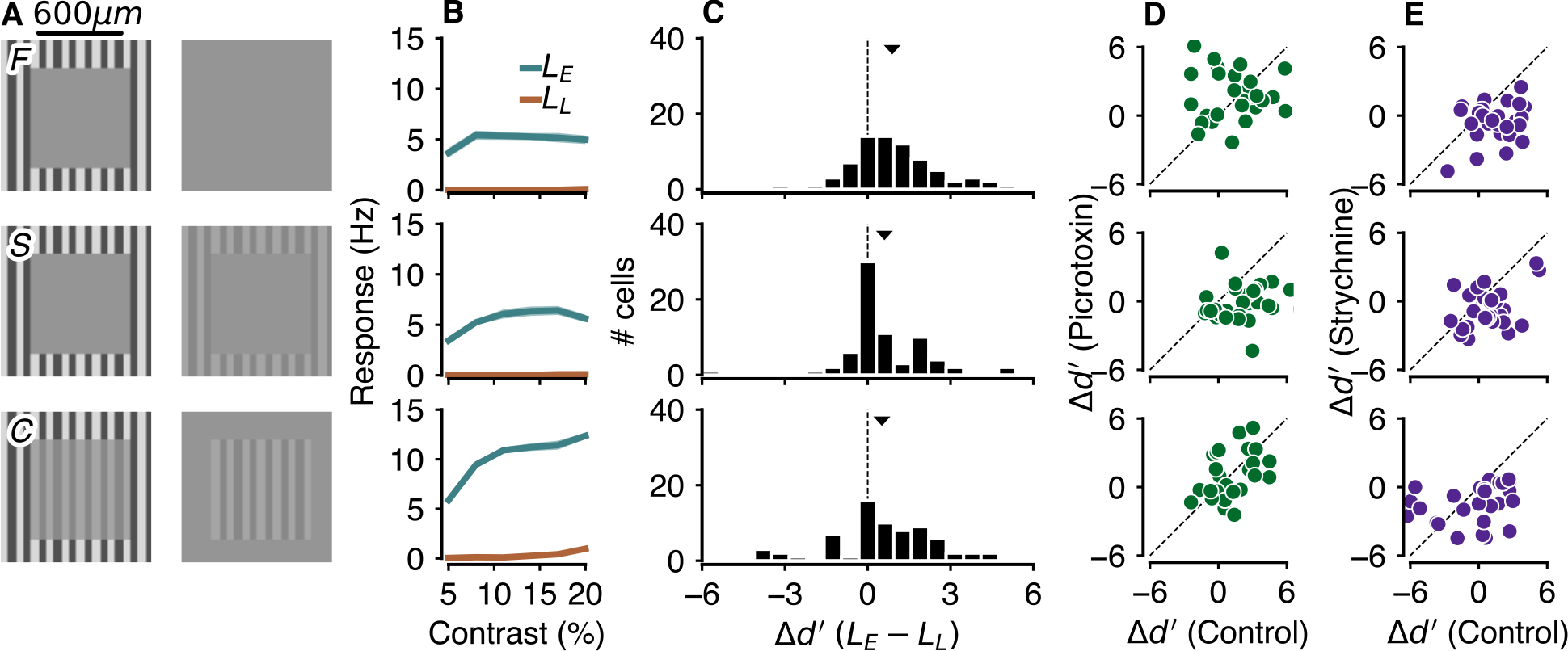
Long range sensitization requires narrow- and wide-field amacrine transmission. (A) Spatial stimulus during adapter (left column) and probe (right), used to probe the long-range sensitization circuit. The stimulus consisted of three conditions, which separately probed changes in discriminability in the center and surround stimulus regions. In the Full-field condition (top row), the stimulus probed both center and surround regions, identically to Fig. 1. The Surround-only condition (middle row) stimulated the surround region only. During the Center-only condition (bottom row), only the center region was stimulated during the probe. Note that in all three conditions, sensitization was triggered by the change in contrast in the surround region, while either the center, surround, or both regions were stimulated during the probe. (B) Response curves for one example cell during each of the three stimulus conditions. Lines show the average response across trials during *L*_*early*_ (green) and *L*_*late*_ (orange). Shaded areas show ± 1 s.e.m and are obscured by data lines. (C) Distribution of change in discriminability between *L*_*early*_ and *L*_*late*_ for 72 RGCs. Arrows indicate population means, which are significant for each stimulus condition. Means ± 1 s.e.m are: 1.03 ± 0.17 (p = 1.08^−7^) for Full-field condition, 0.58 ± 0.17 (p = 0.0014) for Surround-only condition, and 0.68 ± 0.28 (p = 0.016) for Center-only condition. (D-E) Effect of the application of either picrotoxin (D) or strychnine (E) on the change in discriminability between *L*_*early*_ and *L*_*late*_ for 27 ganglion cells, in different preparations than in panel (C). The change in discriminability Δ*d*′ in control trials is plotted on the ordinate, with Δ*d*′ after application of the agent on the abscissa. Picrotoxin disrupted the effect only during the Surround-only condition (middle row, mean of 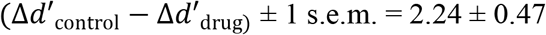, p = 7.093^−5^), while the other two conditions are unaffected (Full-field: −0.18 ± 0.65, p = 0.785; Center-only: 0.58 ± 0.59, p = 0.338). Strychnine disrupts the effects in all conditions (Full-field: mean ± 1 s.e.m = 1.94 ± 0.42, p = 9.764^−5^; Surround-only: mean ± 1 s.e.m. = 1.50 ± 0.40, p = 8.101^−4^; Center-only: mean ± 1 s.e.m = 1.04 ± 0.56, p = 0.0796).

Previous work has shown that local sensitization may be explained by a model of parallel excitatory and inhibitory channels ^13^, in which the output of the inhibitory channel modulates the excitatory channel. Though both channels adapt to strong stimulation, more prolonged adaptation of inhibition leads to a period during which excitation outweighs inhibition, resulting in higher sensitivity until inhibition recovers. The general circuit motif of parallel excitation and inhibition is widespread in the retina^15^, and is also thought to support a phenomenon similar to sensitization the somatosensory cortex ^16^.

The large distances relevant for long-range sensitization suggest that wide-field amacrine cells, which in the salamander are glycinergic ^17^, may contribute to the phenomenon. Yet, GABAergic transmission, which in the salamander arises from narrow-field amacrine cells, is necessary for local sensitization ^13^, and thus we tested whether both types of inhibitory neurotransmission were required for long-range sensitization. We repeated the experiments in Fig. 3A, with the addition of 100 μM picrotoxin to block narrow-field amacrine cell transmission ^17^ or 10 μM strychnine to block wide-field amacrine cell transmission.

In the Center-only condition, only strychnine reduced sensitization, consistent with the model that wide-field inhibitory neurons alone deliver adapting inhibition from the peripheral region to influence excitation in the center (Fig. 3D and E, bottom row). In the Surround-only condition, we expected that only picrotoxin would reduce sensitization because the adapting and probe stimuli were in the same location, and thus long-range communication might not be needed. However, we found that sensitization was reduced by both picrotoxin and strychnine, indicating that, because the stimulus covers a large area, both narrow-field and wide-field inhibition influence excitation from the surround (Fig. 3D and E, middle row). In the Full-field condition, only strychnine significantly disrupted sensitization (Fig. 3D and E, top row). The Full-field condition is a sum of Center-only and Surround-only conditions, and thus one might expect GABAergic inhibition therefore to be required for some amount of sensitization. Although long-range excitation exists in the retina^18–20^, peripheral excitation has a much weaker effect in the presence of a central stimulus ^18^. Furthermore, because glycinergic inhibition is required for both Center-only and Surround-only, we interpret the lack of significant effect of picrotoxin in the Full-field condition as resulting from that the fact that wide-field inhibition has a proportionally larger effect on this stimulus than narrow-field inhibition.

### A model of long-range sensitization

These results suggest a simple model of the retinal circuit underlying long-range sensitization. Extending the previous model of local sensitization ^13^, a sensitization “module” consists of excitation and both adapting narrow-field and wide-field inhibition. This module is tiled across the retina in a repeating circuit motif, being present in both the central and peripheral regions. The ganglion cell then sums the output from these modules (Fig. 4A).

This model accounts for our pharmacological manipulations. Inactivation of wide-field inhibition disrupts long-range sensitization in the Surround-only stimulus (Fig. 3E, middle row) because excitatory responses to the surround stimulus are influenced by wide-field inhibitory input from more distant regions responding to the same surround stimulus. The same connectivity in the central module explains why wide-field inhibition is necessary in the Center-only stimulus; adaptation of the glycinergic wide-field amacrine cells is triggered by the surround stimulus, but the target of this modulation is the central excitatory pathway. This model also includes narrow-field inhibition in the central module, which is the previous model of local sensitization ^13^, but not tested again here. This model reproduced the major characteristics of long-range sensitization in the Full-field, Center-only, and Surround-only stimulus conditions (Fig. 4B-D), including large increases in firing rate following the reduction in surround contrast, and increase sensitivity (slope of response functions, Fig. 4E-G) for a brief period after this reduction of surround contrast.

**Figure 4.**
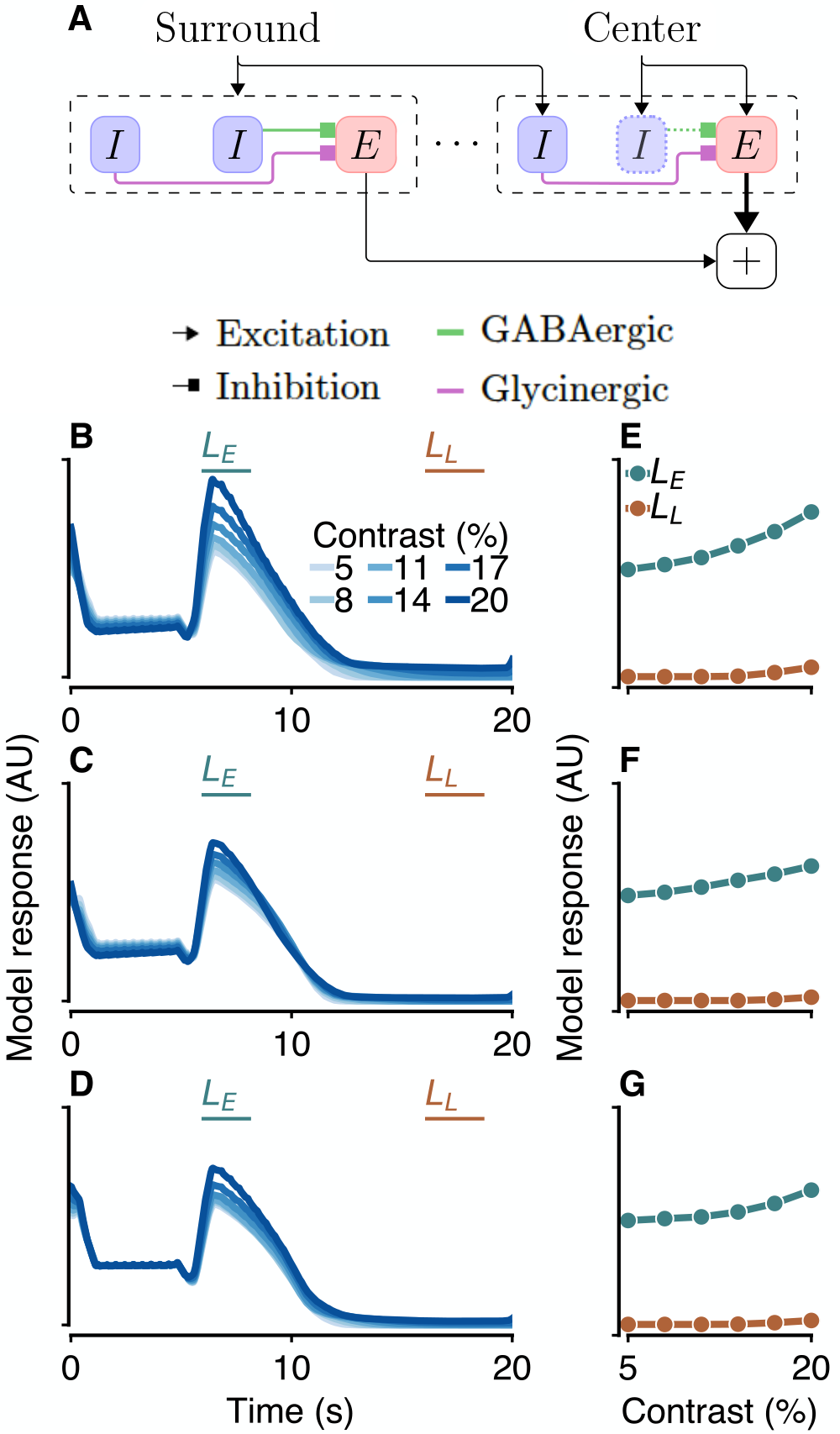
A simple model of long-range sensitization. (A) Schematic of a simple model which reproduces long-range sensitization. The model consists of multiple, tiled “modules” (outlined in dashed black), each of which contains parallel inhibitory (blue) and excitatory (red) channels. The inhibitory channel measures the local stimulus intensity, triggering sensitization, and modulates a potentially distant excitatory response to the stimulus. Both excitation and inhibition contain a linear filter, static nonlinearity, and adaptive stage, with the output of the inhibitory channels combined with the excitatory channel before the excitatory nonlinearity. The two inhibitory channels represent wide-field (purple) or narrow-field (green) inhibition. In the leftmost module, all channels are stimulated by the surround stimulus region (black arrow). In the rightmost module, the excitatory and narrow-field modules are stimulated by the center stimulus, while the wide-field inhibitory channel receives the surround stimulus. (Note that, as sensitization requires a measurement of a changing stimulus, and the center stimulus never changes contrast, the narrow-field channel at right is grayed out.) (B-D) The response of the model to the stimuli in Figure 3. (B) shows the response to the Full-field stimulus condition, (C) to the Surround-only and (D) to the Center-only. Lines above the axes show the early (green) and late (orange) time windows. See Methods for model implementation details. (E-G) Response curves during *L*_*early*_ and *L*_*late*_ for the corresponding stimulus condition in (B-D). In all conditions, the response curves during *L*_*early*_ have both higher mean and higher slope than during *L*_*late*_, indicative of long-range sensitization.

### Long-range sensitization in human perception

Although many complex retinal phenomena have been described ^7^, their relevance to human visual perception in most cases is unknown. To test for the existence of long-range sensitization in human perception, we designed a psychophysical task to parallel our in vitro retinal experiments (Fig. 5A-C). Subjects viewed stimuli on a computer display, fixating on the cross at the right edge of the screen. As in the retinal experiment, the task consisted of a peripheral adapting stimulus, followed by a central probe stimulus. The adapting stimulus was a large Gabor annulus (0.5 cyc/°, s.d. 20°) drifting at 1 cyc/°for 3s, with either high contrast (50% Michelson contrast, Fig. 5A), or low contrast (5%, Fig. 5B). The adapting stimulus then disappeared, and following a brief, variable delay (0, 0.5, or 1s), a small, low contrast probe patch appeared in the center of the annulus for 0.25s (Fig. 5C, 0.5 cyc/°, s.d. 4°). The probe had one of several fixed contrasts, chosen to span each subject’s detection threshold, which was determined by an adaptive procedure in a separate set of trials (see methods for details). Subjects were given a brief response window after the test patch disappeared within which to indicate if they saw the grating. In 10% of all trials the contrast of the probe patch was zero.

We computed the positive response rate for each subject, for each condition (delay interval and test contrast) and fitted sigmoidal psychometric functions (PMFs) to their choice data, which represented their positive choices as a function of probe stimulus contrast (Fig. 5D-F). For all three subjects, the PMFs during trials with the surround adapter were shifted leftward as compared to the baseline PMFs. At an interval of 0 and 0.5s, subjects on average show an increase in the slope of the PMF that was significantly greater than zero (mean ± 1 s.e.m., 10.55 ± 2.99, p=0.024, for 0s interval and 12.72 ± 3.47, p=0.015, for 0.5s) This indicates that the presence of the surround grating increased subjects’ sensitivity to the low-contrast central grating during the early portions of the probe period. This increased sensitivity immediately following the disappearance of the surrounding stimulus mirrors the time course of sensitivity changes in the retinal data. Although we cannot conclude from this experiment that the perceptual effect arises in the retina, this demonstrates that long-range sensitization as seen in the in vitro retina is a phenomenon relevant to human perception.

**Figure 5.**
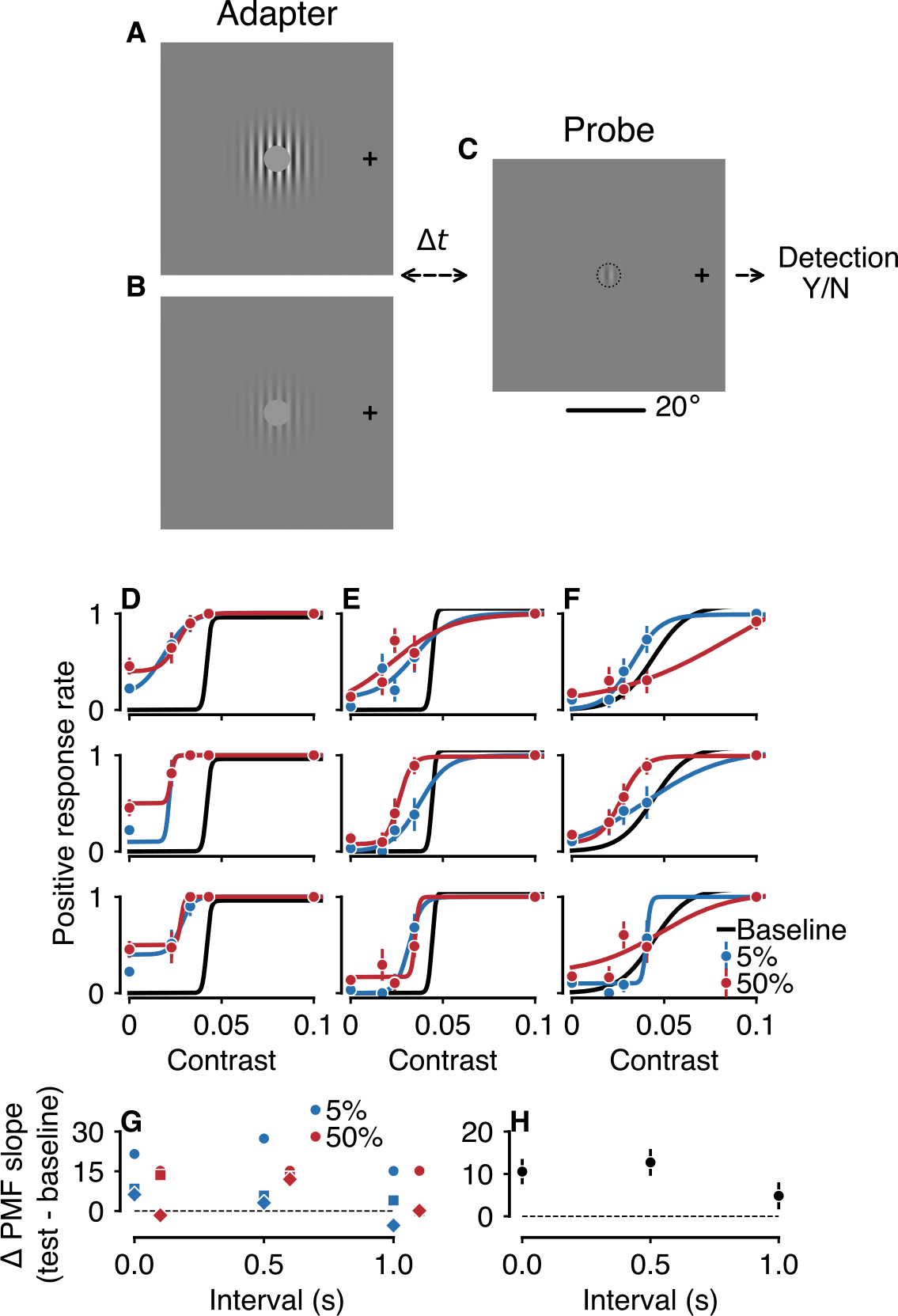
Long-range sensitization in human perception. (A-C) Task setup. While fixating on a spot, three subjects were shown a large Gabor annulus (s.d. = 20°) in the surround, with either 50% (A) or 5% (B) contrast. The surround grating drifted at 1 cyc/° for 3 seconds, after which there was a variable delay (Δ*t* = 0, 0.5, or 1s) during which the screen was blank. Following the delay, a small low (possibly zero) contrast probe Gabor (s.d. = 4°) appeared inside the inner ring of the surround annulus (C, small dotted line indicates surround inner diameter for illustration). Both gratings had a spatial period of 1°. The probe grating appeared for 0.25s, after which the subject indicated whether they detected the grating by pressing a key. No feedback was given. (D-F) Response rates and psychometric functions (PMFs) for each subject, with each of the three intervals shown in the top, middle, and bottom rows, respectively. Black curves show baseline PMFs from threshold-estimation trials, which describes each subject’s detection performance as a function of contrast without the surround adapter. Blue points and lines show each subject’s choice data and fitted PMFs during trials with the 5% contrast surround grating, and red during trials with the 50% contrast surround grating. PMFs during trials in which the surround was present are shifted leftward relative to baseline for all subjects. (G) The average slope difference between test and baseline PMFs of each curve from D-F, plotted as a function of the interval between adapter and probe. Colors indicate the contrast of the surround, matching the PMF from which they are derived, with each symbol marking a single subject. (H) The average change in sensitivity between test and baseline, averaged across all subjects and both adapter contrasts (error bars show ± 1 s.e.m.). Sensitivity is significantly higher during the test (mean ± 1 s.e.m., 10.55 ± 3.28, p=0.024, and 12.72 ± 3.47, p=0.015, for 0s and 0.5s, respectively).

## Discussion

Here we have shown that strong surround stimulation, outside the receptive field center of ganglion cells drives a prolonged change in the responses to stimuli presented in the center. After a reduction in surround contrast, ganglion cells display increases in firing rate, as well as increased sensitivity to contrast in the center (Fig. 1). This increased sensitivity improves the ability of ganglion cells to discriminate local changes in stimulus contrast. Long-range sensitization also exhibits a form of pattern-specificity. In stimulus conditions in which the center and surround regions share spatial frequency statistics, ganglion cells sensitize more than when the statistics of the two regions differ (Fig. 2).

The model proposed here naturally translates to a circuit description of sensitization, with excitation from the center provided by bipolar cells, and adapting inhibition for local stimuli provided by narrow-field amacrine cells ^13^ and for more distant stimuli delivered by wide-field amacrine cells. One circuit element that is not clear is the source of excitation from the peripheral stimulus. Although peripheral excitation has been shown ^18–20^the neurons that convey this excitation are unknown. In principle, excitation could result from disinhibition through serial inhibitory connections, but our finding that neither strychnine nor picrotoxin eliminate excitation from the peripheral stimulus (Fig. 3) argues against this.

The response properties of wide-field amacrine cells provide a possible explanation of the observed pattern-specificity: the nonlinear subunits of wide-field amacrine cells are strongly driven by the high spatial-frequency gratings used throughout these experiments ^21^. Wide-field amacrine cells are a broad and diverse class of interneurons, whose complete response properties are far from known. One might expect that a full understanding of these properties would provide additional information as to the pattern specificity of sensitization.

It is known that adaptation within the receptive field center is pattern specific to a set of stimulus correlations that include spatial frequency, orientation, and temporal correlations ^22^. On the other hand, the pattern specificity of *local* sensitization is not known. While local sensitization is triggered by both first- and second-order features (luminance and contrast) ^12^, it is not clear if this phenomenon increases sensitivity specifically to the triggering statistic, as it does in long-range sensitization.

Based on our results, one can propose a general neural circuit that could exhibit sensitization to the stimulus statistics or features to which the inhibitory channel is sensitive. This provides a simple but powerful mechanism by which to understand prediction in neural systems. A large body of work characterizes complex neural processes, especially perception^23–26^, as one of inference, in which the current sensory evidence is combined with prior expectations. Yet, the exact neural mechanisms through which such inference occurs are unclear. Adaptation, among the best-studied dynamic properties of neural systems, predicts that repeated or strong presentation of a stimulus feature should lead to a decrease in the responsiveness to that feature, yet inference requires such stimulation increase the system’s responsiveness.

Sensitization provides a general mechanism through which such increases may arise, using the parallel circuit described here. Given an inhibitory pathway sensitive to a specific stimulus feature, adaptation of the modulatory transmission of that pathway onto the parallel excitatory pathway could generate an inference that this feature is more likely. The only requirements for such a general circuit mechanism are the existence of inhibitory and excitatory pathways that are sensitive to the same feature, both of which are loose requirements. Such a circuit could be used to drive inference about arbitrary stimulus features, providing a neural basis for the complex inference processes observed throughout human perception.

Human perception is furthermore often characterized as following the more specific process of Bayesian inference. This process too may be captured by the sensitization circuit proposed here. The gain of the inhibitory channel transmission could effectively encode the inverse probability of its predominant feature; thus, halving the gain implies a doubling of the feature’s prior probability. This assumes that the outputs of the inhibitory and excitatory channels are multiplied, and that activity in the excitatory channel represents the likelihood of the visual feature from the direct sensory input. As before, these requirements are fairly mild, and in fact, local retinal sensitization caused by a small object conforms to such a Bayesian inference model^13^.

Here, we have provided evidence for an extension of the sensitization circuit, which encompasses multiple retinal cell types and visual features. This suggests that the underlying mechanism are broad and provide the basis for a simple neural circuit with sufficient flexibility to implement the inference processes that describe human perception.

## Materials and methods

### Experimental preparation

To record the simultaneous spiking activity of multiple ganglion cells, retinae from larval tiger salamanders were isolated under dim red light beneath a microscope fitted with infrared eyepieces. Retinae were bathed in oxygenated Ringer’s solution (110 μM NaCl, 5 μM KCl, 1 μM CaCl_2_, 1.6 μM MgCl_2_, 10 μM D-glucose, Sigma-Aldrich) at 20-22°C, buffered with 22 μM NaHCO_3_. Electrical activity was recorded on an array of 60 microelectrodes (MEA, Multichannel Systems), acquired and visualized in real-time using custom software written in C++. Action potentials from individual ganglion cells were isolated using custom spike-sorting software, written in MATLAB, C++ and Python. Each retina resulted in ~20-30 isolated ganglion cells and results here combine across at least 2 retinas for each experiment. Numbers of cells are quoted in each individual experiment when relevant.

### Visual stimulation

Stimuli were drawn and presented using MATLAB and the Psychophysics Toolbox, on a linearized CRT monitor with a mean power of 10 m*W*/*m*^2^ and an effective frame rate of 30 Hz. Stimuli were optically reduced through a microscope objective lens (Nikon Plan Fluor 10x, WD 3.5cm) for presentation onto the retina. Checkerboard patterns used to measure receptive fields were generated by drawing the luminance of each checker (100μm on each side) on each frame from a Gaussian distribution with mean μ, and standard deviation σ. In this case, contrast was defined as *c* = *μ*/*σ* with *μ* given by the gray level of the monitor. Other stimuli use Michelson contrast, defined as (*I*_*max*_ - *I*_*min*_)/(*I*_*max*_ + *I*_*min*_), where *I*_*max*_ and *I*_*min*_ are the maximal and minimal monitor luminance values, respectively.

### Contrast response curves

For each contrast, *c*, the mean firing rate *m*(*c*) over a time interval *T* was computed for each response, *r*_*c*_ as

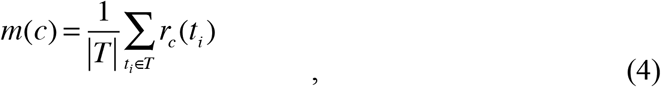

for time intervals T = *L*_*eariy*_ or *L*_*late*_.

### Linear-nonlinear models

To constrain the model described below, we estimated linear-nonlinear models for some cells from their responses to spatiotemporal white noise stimulation. Receptive fields were estimated via reverse-correlation, using the standard method of spike-triggered averaging (STA) ^27^. Given the time-varying stimulus *s*(*t*, *x*, *y*) with *x* and *y* indicating spatial dimensions, the filter for each cell was computed as

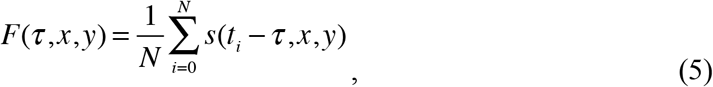

where *s* is the mean-subtracted stimulus, and *τ* is the time preceding a spike at time *t*_*i*_. The filter was then normalized in amplitude so that when convolved with the stimulus it left the stimulus variance unchanged ^28^.

A static nonlinearity for each cell was estimated by comparing the linear prediction from the above linear filter with the true firing rate. The linear prediction *g* was estimated as the convolution of the stimulus and filter:

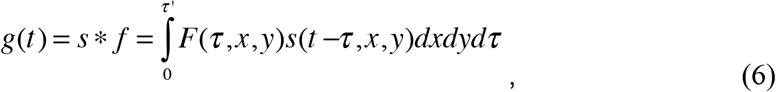

where *τ*′ is the duration of the filter. The nonlinearity *N*(*g*) was then computed as the expected value of the firing rate given each value of the predicted rate

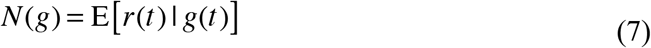

by sorting the linear prediction *g*(*t*) and the true firing rate *r*(*t*) using *g*(*t*) as the key. The prediction, *g*(*t*) was binned into groups of nearly-equal size, and *r*(*t*)averaged within each bin. This defined a piecewise-linear function mapping predicted to true rate. All methods for constructing LN models were implemented in the Pyret software package^29^.

### Pharmacology

Pharmacological manipulations were performed to explore the retinal mechanisms underlying long-range sensitization in ganglion cells. These experiments incorporated either 100 μM picrotoxin or 20 μM strychnine (Sigma-Aldrich) into the standard Ringer’s solution described above. The solution with each agent was washed in after some number of control blocks, and analyses were performed separately on control and experimental blocks. The first block after the agent was washed in were ignored (≈ 1000s).

### Model

To model the phenomenon of long-range sensitization, we combined two sensitization modules ^12^, a “surround module”, *M*_*S*_, with excitation and inhibition both deriving from the surround, and a “cross module”, *M*_*C*_, with wide-field inhibition arising from the surround, and narrow-field inhibition and excitation arising in the center. Each module contains an inhibitory and excitatory channel, connected by a synapse. Each channel consists of a linear filter, nonlinearity, and adaptive block, implemented as divisive feedforward inhibition. The output of the inhibitory channel was combined with the excitatory channel prior to its nonlinearity. In the surround module, *M*_*S*_, both the excitatory and inhibitory channels were driven by the background stimulus. In the cross module, *M*_*C*_, the inhibitory channel was driven by the background stimulus, and the excitatory channel by the center.

Each module was implemented as follows. Linear filters and nonlinearities were estimated from a representative cell’s LN model in response to white noise stimulation. The stimulus s was convolved with each linear filter as in (Eq. 6). The output of the inhibitory linear filter *l* was then passed through the nonlinearity estimated from data. This output, *n*, was passed through a feedforward, divisive inhibition stage, implemented as:

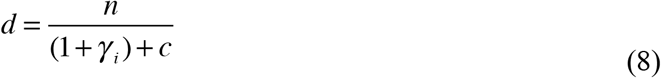

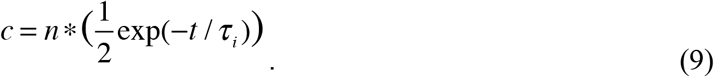

The inhibitory channel output was convolved with a filter modeling synaptic delay,

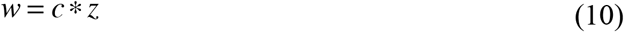

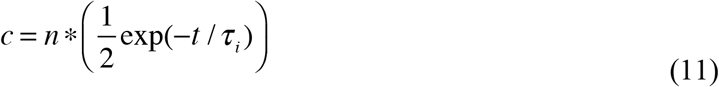

and passed through a sigmoid nonlinearity:

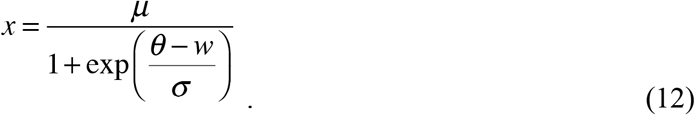

The output of the synapse was combined linearly with the output of the excitatory linear filter, and passed through a similar nonlinearity and adaptive block, followed by a final rectifier nonlinearity,

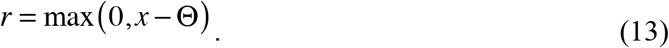

The parameters for the surround module (*M*_*S*_) and the cross module (*M*_*C*_) were as follows:

| | $\tau_i$ | $\gamma_i$ | $\alpha$ | $\mu$ | $\theta$ | $\sigma$ | $\tau_e$ | $\gamma_e$ | $\Theta$ |
| --- | --- | --- | --- | --- | --- | --- | --- | --- | --- |
| $M_S$ | 3.0 | 1.0 | 0.15 | $5.3 \times 10^{-5}$ | 0.7 | 0.05 | 3.0 | 1.0 | 0.5 |
| $M_C$ | 2.5 | 0.7 | 0.15 | $5.3 \times 10^{-5}$ | 0.6 | 0.025 | 5.0 | 1.0 | 0.725 |

### Psychophysics

To assess the perceptual relevance of long-range sensitization in the retina, human subjects (an author and two naive subjects) performed a simple psychophysical task. All methods and procedures were approved by the Stanford Institutional Review Board. Subjects viewed the stimulus stereoscopically, through a polarized beam-splitter and polarized glasses (Planar, monitors synchronized with Matrox DualHead2Go). During the task, subjects fixated at the right edge of the monitors, and stimuli were presented at 15° eccentricity. Subjects were shown brief presentations of drifting Gabor patches, and asked to indicate when they detected a grating by pressing a key. No feedback was given, and if the subject did not respond within an allowed time window, the next trial began. Subjects performed an initial set of trials to measure detection thresholds, using an adaptive staircase procedure (QUEST^30^). Subjects’ thresholds were used to determine an appropriate range for experimental stimuli, which used the method of constant stimuli to measure the proportion of positive responses as a function of contrast. Psychometric functions were fitted as function of the contrast *c* to the response rate for each subject and condition, using a sigmoid function:

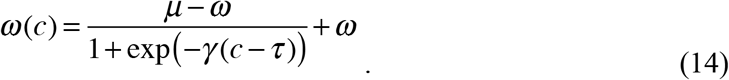

See Results for details of the stimulus and task design.

## Acknowledgements

The authors wish to acknowledge Anthony Norcia for helpful discussions and for the use of equipment for psychophysical stimulus presentation, and Surya Ganguli and Eric Knudsen for helpful discussions. This work was supported by grants from the NEI (SAB) and NRSA EY025110 (BN).

## Author contributions

B.N. and S.A.B. designed the study, B.N. performed the experiments, B.N. and S.A.B. developed the analyses, and B.N. and S.A.B. wrote the manuscript.

